# Resolving the *CES1* Genomic Locus with Cas9-Directed Targeted Long-Read Sequencing for Precision Pharmacogenomics

**DOI:** 10.64898/2025.12.03.692062

**Authors:** E. Lekka, A. Ambrodji, A. Nater, A. Ballah, U. Amstutz, A. Ramette, C.R. Largiadèr

## Abstract

Carboxylesterase 1 (CES1) is the primary hepatic hydrolase in humans, crucial for the metabolism of ester-containing drugs and endogenous lipids. However, the *CES1* genomic region is difficult to resolve because of adjacent highly homologous pseudogenes and the presence of large structural variants. These complexities often cause read misalignment and inaccurate variant calling with conventional short-read sequencing, hindering reliable pharmacogenomic analyses. To overcome these limitations, we employed an optimized, PCR-free Oxford Nanopore Technologies (ONT) sequencing method, Cas9directed targeted sequencing (nCATS), to characterize the targeted region of up to 76 kb, including *CES1, CES1P1* or *CES1A2*, and their intergenic regions. This approach uses Cas9 to selectively enrich and sequence long native DNA fragments, while avoiding amplification-induced artefacts. Long-read sequencing was performed in 23 human blood samples and the HepG2 hepatoblastoma cell line enabling high-resolution mapping to the *CES1* locus. We uncover five previously unrecognized main *CES1* haplotypes and report that many single nucleotide variants (SNVs) in public databases are likely artefacts caused by short-read misalignment. Additionally, we identify long inverted repeats (LIRs) flanking a fragile genomic site within the region, which may form DNA hairpins and contribute to structural plasticity at the locus. This study demonstrates the utility of long-read sequencing for resolving complex genomic regions such as *CES1*, allowing comprehensive detection of structural variants and haplotype-resolved SNVs. Our findings provide improved reference sequences and deeper insight into *CES1* diversity, with significant implications for future pharmacogenetic research and the development of personalized treatment strategies involving CES1-metabolized medications.

## INTRODUCTION

Carboxylesterase 1 (CES1) is a serine esterase highly expressed in the liver and other metabolically active tissues. It plays a critical role in the metabolism of ester- and amide-containing drugs (Imai et al. 2006; Rasmussen et al. 2015; Her and Zhu 2020)(https://www.pharmgkb.org/). In addition to xenobiotics, CES1 is involved in lipid metabolism (Lian et al. 2018; Li et al. 2023). *CES1* has emerged as an important pharmacogene due to its broad substrate specificity, which includes diverse therapeutic agents such as anticancer prodrugs (Longley et al. 2003), opioids, stimulants (McRee 2003), angiotensin-converting enzyme inhibitors (Wang et al. 2016), anti-platelet agents (Zhu et al. 2013), and antiviral medications against influenza (Zhu and Markowitz 2013) or COVID-19 (Li et al. 2021). Given its influence on the pharmacokinetics and safety profiles of many clinically relevant compounds, comprehensive characterization of the *CES1* genetic locus may yield important insights and support the development of genotype-guided therapeutic strategies.

Sequencing of the *CES1* locus presents considerable challenges due to its repetitive structure and genomic complexity. Located on chromosome 16, *CES1* spans approximately 30 kb, comprises 14 exons, and lies adjacent to two homologous pseudogenes, *CES1P1* and *CES1P2* (Fukami et al. 2008). *CES1* is subject to structural variation and several haplotypes have been reported (Fukami et al. 2008; Hosokawa et al. 2008). In one common haplotype, *CES1P1* (consisting of six exons) is arranged in a tail-to-tail orientation with the functional *CES1* gene (also referred to as *CES1A1*). In another haplotype, *CES1P1* is replaced by *CES1A2* (also known as *CES1P1VAR)*, an inverted duplication of *CES1* (Tanimoto et al. 2007). Sequence differences in the promoter between *CES1A1* and *CES1A2* lead to significantly lower transcription efficiency of *CES1A2* (Fukami et al. 2008; Hosokawa et al. 2008). However, some individuals carry a promoter haplotype of *CES1A2* with two overlapping binding sites for the transcription factor Sp1, resulting in significantly increased *CES1A2* transcription (Yoshimura et al. 2008). There are only a few nucleotide differences in exon 1 between *CES1A1* and *CES1A2*, which result in four amino acid substitutions in the open reading frame, producing distinct protein isoforms. Exon 1 encodes a signal peptide (Shibata et al. 1993) which contributes to the subcellular localization differences observed between the isoforms: CES1A1 predominantly localizes to the endoplasmic reticulum, while CES1A2 is distributed in both the endoplasmic reticulum and cytosol (Satoh and Hosokawa 2006). Three subtypes of *CES1A1* (*CES1A1a, CES1A1b, CES1A1c*) have been reported, and both *CES1A1b* and *CES1A1c* produce *CES1A2* type of mRNA (Tanimoto et al. 2007).

Numerous studies have examined the impact of *CES1* genetic variation on the metabolism of its substrate drugs (Zhu et al. 2008; Sai et al. 2010; Wang et al. 2016; Hamzic et al. 2017; Stage et al. 2017a). Both structural variations and single nucleotide variants (SNVs) have been shown to influence CES1 enzymatic activity, thereby affecting the efficacy and/or toxicity of CES1-metabolized compounds. Missense variants may alter the CES1 catalytic function by changing the structural environment of the active site or by disrupting protein folding and stability (Zhu et al. 2008; Zhu and Markowitz 2009; Hussain et al. 2024) while SNVs that reside on *CES1* promoter can alter transcriptional activity (Geshi et al. 2005). One well-characterized functional variant is the nonsynonymous SNV rs71647871 (p. Gly143Glu or G143E), which causes a non-conservative substitution of glycine to glutamic acid and has been associated with markedly decreased enzymatic activity (Zhu et al. 2008).

Duplicated loci, such as those found at the *CES1* gene locus, represent some of the most challenging regions of the human reference genome to resolve (Vollger et al. 2022). Because of high sequence similarity among paralogous segments—including *CES1*, *CES1A2*, and *CES1P1*—traditional short-read sequencing often misassigns sequence variants, leading to false identification of SNVs due to read misalignment. As highlighted by Ferrero-Miliani and colleagues, many SNVs currently attributed to *CES1* may in fact originate from *CES1A2* or *CES1P1*, and vice versa (Ferrero-Miliani et al. 2017). This lack of specificity compromises the accuracy of genotyping assays and hinders reliable genotypephenotype associations, particularly in pharmacogenomics studies linking *CES1* variants to drug metabolism and toxicity (Rasmussen and Madsen 2018). To address these challenges, we applied a PCR-free nanopore Cas9-targeted sequencing (nCATS) strategy (Gilpatrick et al. 2020; Watson et al. 2020; McDonald et al. 2021; Alfano et al. 2022; Skowronek et al. 2022; Turner et al. 2023; Iyer et al. 2024; Rausch et al. 2025). This approach exploits the programmable DNA cleavage activity of Cas9 to direct adapter ligation specifically to the genomic interval encompassing *CES1P1* or *CES1A2*, *CES1* and their intergenic region. PCR is associated with amplification bias and chimera formation (Ambrodji et al. 2024). In contrast, this method avoided such PCR artefacts, improved read mappability, and enabled accurate detection and phasing of both structural and small variants.

In our nCATS-based sequencing of 23 genetically diverse human blood samples and HepG2 cells, we identified two distinct *CES1P1* haplotypes and revealed a recurrent inversion within the *CES1P1/CES1A2-CES1* intergenic region. Notably, we identified the presence of long inverted repeats (LIRs) at the *CES1* locus, which may function as precursors to small interfering RNAs (siRNAs), and potentially contribute to the genomic plasticity observed in this region. Finally, leveraging the longread data generated in this study, we re-examine major *CES1* haplotypes and present refined reference sequences for this complex genomic locus.

Overall, our approach combines targeted Cas9-based enrichment with long-read nanopore sequencing, and provides a robust framework for resolving the challenging *CES1* locus with high accuracy. This method not only enhances the characterization of *CES1* genetic variation but also holds promise for enabling more precise genotype-based personalization of therapies involving *CES1*metabolized drugs. Furthermore, the data generated through this strategy provide a valuable resource for investigating the regulatory landscape and functional roles of *CES1* in both physiology and disease.

## RESULTS

### Cas9-targeted nanopore sequencing for comprehensive genetic analysis of *CES1*

Here, we applied nanopore Cas9-targeted sequencing (nCATS) to resolve complex genetic variation, including large structural variants (SVs) at the pharmacogene *CES1*. There are two main structural haplotypes of *CES1*, containing *CES1P1* or *CES1A2* (**Fig. 1A**).

**Figure 1.**
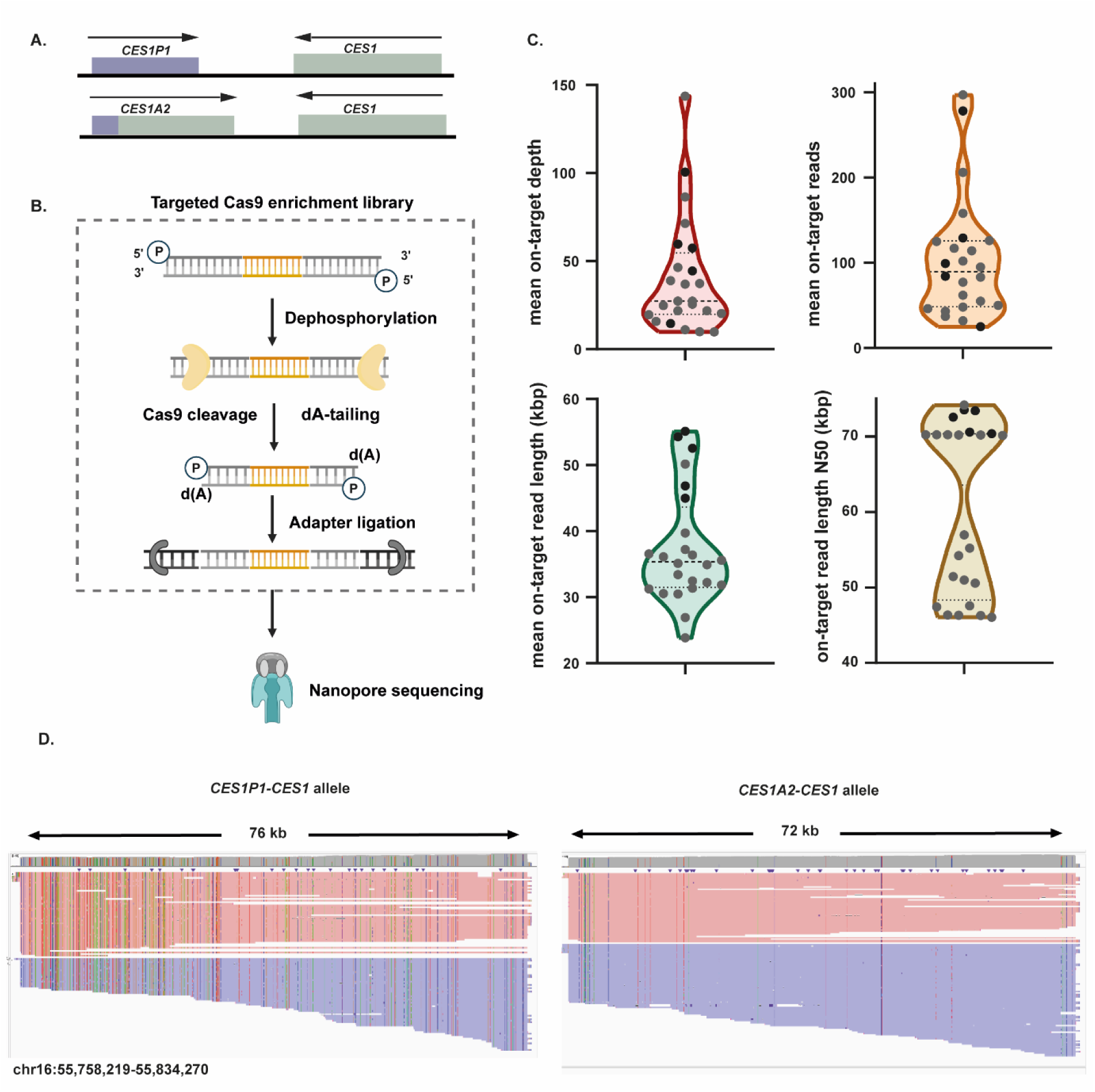
Targeted sequencing of *CES1* locus using nanopore Cas9-targeted sequencing (nCATS). (A) Schematic representation of the two major structural haplotypes at the *CES1* locus, featuring either *CES1P1* or *CES1A2* arranged in a tail-to-tail orientation with *CES1.* (B) Overview of the library preparation workflow used for targeted enrichment of *CES1* using nCATS. (C) Sequencing performance metrics from 23 human blood samples and HepG2 cells enriched for *CES1* using nCATS. Shown are average on-target depth, number of on-target reads, mean on-target read length, and read length N50. Black dots represent samples sequenced with R9.4.1 flow cells; grey dots represent samples sequenced with R10.4.1 flow cells. (D) Example of an nCATS-sequenced *CES1P1/CES1A2* heterozygous sample aligned to a custom two-contig haplotype reference containing the sequences of the two major *CES1* structural haplotypes (*CES1P1–CES1* and *CES1A2–CES1*). Aligned reads were visualized using Integrative Genomics Viewer (IGV).

A critical component of the nCATS protocol is the design of guide RNAs (gRNAs), which determine Cas9 cleavage sites and direct the ligation of sequencing adapters to target regions. To enrich for the *CES1* locus, we initially compared two Cas9 cleavage strategies using the same genomic DNA sample (**Supplemental Fig. S1**). The first approach, referred to as the *multi-target cleavage scheme*, involved generating separate libraries for *CES1P1* or *CES1A2* and *CES1*, which were later pooled before loading onto the nanopore flow cell. For *CES1P1*, a double-cut excision strategy was employed using two gRNAs flanking the locus. In contrast, *CES1A2*, when present, was cleaved only at the upstream boundary. Because *CES1* spans approximately 30 kb, we employed a bricklayer tiling approach, recommended by Oxford Nanopore Technologies (ONT) for capturing regions >20 kb. For this target, we combined tiling with excision to increase the likelihood of obtaining full-length reads across *CES1*.

In the second approach, designated as the *end-to-end cleavage scheme*, we targeted a broader genomic region encompassing upstream *CES1P1* or *CES1A2* through to upstream *CES1* using two gRNAs on each flank. This strategy was designed to capture long reads across the entire region of interest. Before DNA library preparation, gRNA efficiency was validated using PCR-amplified Cas9-RNP substrates **(Supplemental Figs. S2-S4).**

Both gRNA strategies generated reads spanning the full target region; however, the end-to-end approach yielded longer reads and higher coverage (**Table 1**). This method was also more costeffective, reducing the number of reaction steps.

**Table 1.** Alignment metrics for the multi-target and end-to-end sequencing runs of the same *CES1P1/CES1A2* heterozygous human blood sample S1 with R9 nanopore chemistry.

| Cleavage Scheme | Target | On-target Reads | Mean on-target read length (bp) | Read length N50 (bp) | Covered bases (%) | Mean target sequencing depth |
| --- | --- | --- | --- | --- | --- | --- |
| Multi-target | *CES1P1-CES1* | 219 | 24,351 | 34,255 | 76,047<br>99.99 | 68.44 |
| Multi-target | *CES1A2-CES1* | 235 | 23,813 | 33,674 | 72,741<br>99.99 | 75.01 |
| End-to-end | *CES1P1-CES1* | 132 | 53,519 | 73,194 | 76,028<br>99.97 | 90.03 |
| End-to-end | *CES1A2-CES1* | 146 | 56,582 | 70,150 | 72,745<br>100.00 | 111.47 |

Based on these findings, we selected the end-to-end scheme for all subsequent library preparations. This strategy enabled single-molecule nanopore sequencing of long genomic segments (>70 kb), capturing upstream *CES1P1* or *CES1A2*, the intergenic region, the *CES1* gene, and a portion of its upstream regulatory elements. As such, it enabled comprehensive detection and phasing of structural and small variants without the need for read assembly.

We initially screened 146 human blood samples from the Liquid Biobank Bern (https://www.biobankbern.ch/home-en/) for *CES1P1* or *CES1A2* using allele-specific long-range PCR (**Supplemental Fig. S5, S6**). Bidirectional Sanger sequencing was then performed to identify the *CES1A1a, CES1A1b,* and *CES1A1c* genotypes as well as the 143E allele. We observed substantial diversity at the *CES1* locus in the analyzed biobank samples, reflected by a wide range of haplotypes (**Supplemental Fig. S6**). In the study population, the allele frequency of the haplotype carrying *CES1A2* was 21.91%. A total of 42 individuals (28.76%) carried one copy of *CES1A2*, and 11 individuals (7.5%) carried two copies. The *CES1A1c* subtype had an allele frequency of 17.12%, with 32 heterozygous carriers (21.91%) and nine homozygous individuals (6.16%). In contrast, the *CES1A1b* subtype was rare, with an allele frequency of 0.34% and only one heterozygous carrier (0.68%). The 143E allele (rs71647871) was detected in two heterozygous individuals (allele frequency: 0.68%).

We selected a subset of 23 human blood samples representing diverse *CES1* haplotypes for subsequent nCATS analysis (**Supplemental Table S1**). Initially, we applied the nCATS method (**Fig. 1B**) to sequence four human blood samples and HepG2 cells using ONT R9 chemistry, available at that time. Subsequently, we adapted our library preparation protocol for the latest ONT R10 chemistry and sequenced an additional nineteen samples. Sequencing with R9 and R10 chemistries yielded comparable performance metrics (**Fig. 1C**). Across all samples, we achieved an average target coverage of 40.54× (SD = 31.90), with a mean of 103.66 (SD = 69.98) on-target reads, a mean on-target read length of 37.49 kb (SD=8.46), and a mean on-target read length N50 of 60.63 kb (SD = 11.05) (**Fig. 1C**; **Tables 2, 3**).

Structural variation in the *CES1* locus has traditionally been inferred using PCR-based assays, primarily by estimating the *CES1P1* or *CES1A2* copy number (Stage et al. 2017a; Bjerre et al. 2018; Ikonnikova et al. 2022). While informative, these methods are limited in their ability to resolve complex structural configurations and do not directly phase variants. Using our nCATS-based approach, structural haplotypes were inferred by competitively mapping long reads to reference sequences representing the two major configurations at this locus: *CES1P1–CES1* and *CES1A2–CES1* (**Fig. 1D**). All structural haplotypes inferred from nCATS data were concordant with assignments from *CES1P1-* and *CES1A2*specific long-range PCR in the same samples (**Supplemental Table S1**).

**Table 2.**
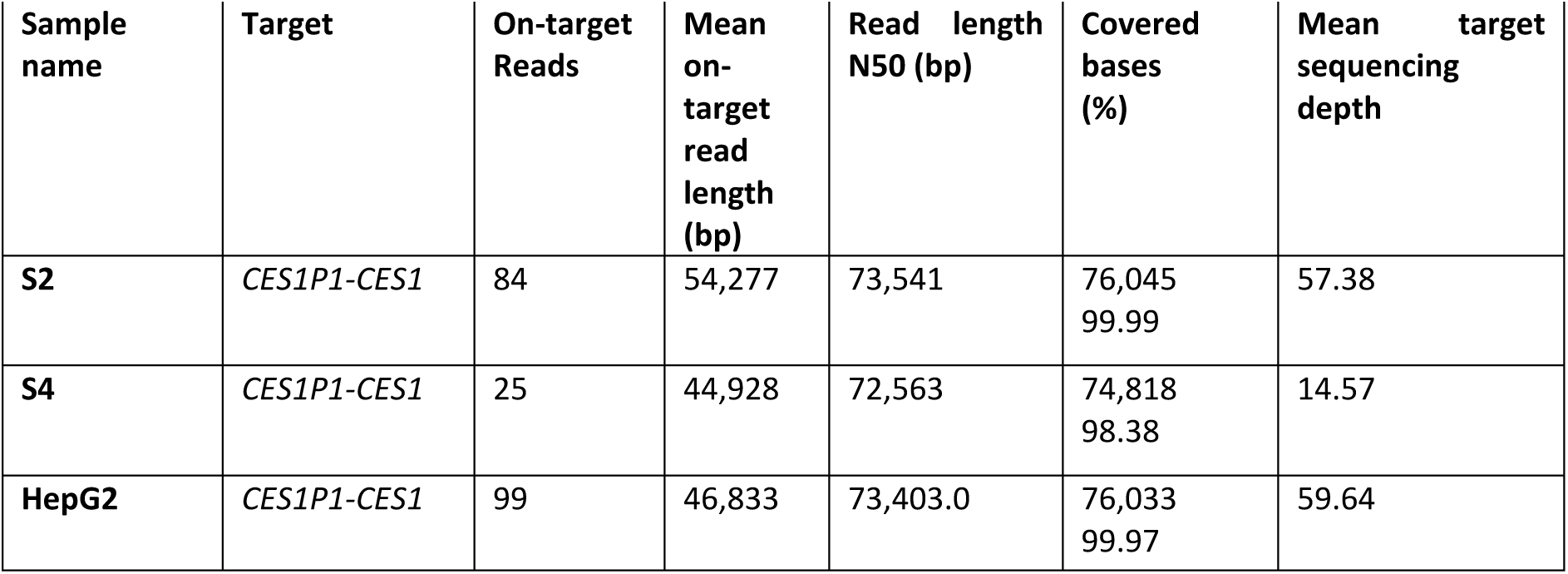
Alignment metrics for human blood samples analysed with R9 nanopore chemistry.

**Table 3.** Alignment metrics for human blood samples analysed with R10 nanopore chemistry.

| Sample name | Target | On-target Reads | Mean on-target read length (bp) | Read length N50 (bp) | Covered bases (%) | Mean target sequencing depth |
| --- | --- | --- | --- | --- | --- | --- |
| **S5** | *CES1P1-CES1* | 74 | 21,978 | 61,271 | 74,831<br>98.39 | 21.27 |
| **S5** | *CES1A2-CES1* | 51 | 32,557 | 46,336 | 72,711<br>99.95 | 22.45 |
| **S6** | *CES1P1-CES1* | 40 | 33,340 | 70,182 | 74,822<br>98.38 | 17.46 |
| **S6** | *CES1A2-CES1* | 62 | 38,628 | 46,336 | 72,744<br>99.99 | 32.56 |
| **S7** | *CES1P1-CES1* | 50 | 31,262 | 70,244 | 74,832<br>98.38 | 20.36 |
| **S8** | *CES1A2-CES1* | 37 | 31,349 | 46,043 | 71,530<br>98.33 | 15.84 |
| **S9** | *CES1P1-CES1* | 62 | 50,137 | 74,162 | 76,002<br>99.93 | 37.03 |
| **S10** | *CES1P1-CES1* | 31 | 39,260 | 68,480 | 74,832<br>98.39 | 15.42 |
| **S10** | *CES1A2-CES1* | 17 | 27,573 | 37,261 | 72,072<br>99.07 | 6.43 |
| **S11** | *CES1P1-CES1* | 85 | 27,524 | 70,183 | 75,998<br>99.93 | 30.28 |
| **S11** | *CES1A2-CES1* | 41 | 36,809 | 45,706 | 72,705<br>99.94 | 20.57 |
| **S12** | *CES1A2-CES1* | 55 | 36,389 | 46,313 | 72,722<br>99.97 | 27.25 |
| **S13** | *CES1A2-CES1* | 43 | 33,417 | 46,332 | 72,716<br>99.96 | 19.62 |
| **S14** | *CES1P1-CES1* | 206 | 32,465 | 70,238 | 74,838<br>98.410 | 86.46 |
| **S15** | *CES1P1-CES1* | 77 | 37,218 | 70,189 | 76,033<br>99.97 | 36.91 |
| **S16** | *CES1P1-CES1* | 46 | 40,701 | 70,203 | 74,811<br>98.37 | 23.94 |
| **S16** | *CES1A2-CES1* | 37 | 38,464 | 46,323 | 71,530<br>98.33 | 19.55 |
| **S17** | *CES1P1-CES1* | 95 | 31,835 | 70,151 | 74,831<br>98.39 | 38.88 |
| **S18** | *CES1P1-CES1* | 117 | 30,472 | 70,204 | 76,031<br>99.97 | 46.42 |
| **S19** | *CES1P1-CES1* | 33 | 30,905 | 70,173 | 74,804<br>98.36 | 13.30 |
| **S19** | *CES1A2-CES1* | 13 | 34,332 | 46,299 | 70,914<br>97.48 | 6.11 |
| **S20** | *CES1P1-CES1* | 158 | 34,933 | 70,219 | 76,032<br>99.97 | 71.48 |
| **S21** | *CES1A2-CES1* | 298 | 35,616 | 46,282 | 72,745<br>100.00 | 143.66 |
| **S22** | *CES1P1-CES1* | 32 | 23,860 | 50,985 | 74,795<br>98.35 | 9.98 |
| **S23** | *CES1P1-CES1* | 53 | 32,262 | 70,163 | 74,812<br>98.37 | 22.33 |
| **S23** | *CES1A2-CES1* | 61 | 39,404 | 46,277 | 72,723<br>99.97 | 32.53 |

Nanopore sequencers offer a software-controlled enrichment strategy known as adaptive sampling or “Read Until” (Loose et al. 2016; Martin et al. 2022; Terrazos Miani et al. 2024; Ulrich et al. 2024). In this approach, the initial ∼400–600 bases of a DNA molecule are analysed in real time, and reads that do not map to a predefined target reference can be actively rejected by reversing the voltage across the nanopore. This process frees individual pores for the capture of new molecules, potentially enriching for target sequences without additional library preparation steps. We examined if our nCATS method could benefit from adaptive sampling by running the same nCATS library in regular (nCATS) or adaptive sampling (nCATS-AS) mode for 24 hours. Compared to the regular mode, adaptive sampling yielded a moderate increase in on-target reads and mean sequencing depth (**Table 4**).

**Table 4.** Alignment metrics for nCATS library with or without adaptive sampling for the same human blood sample S3 using R9 nanopore chemistry.

| Sample name | Target | On-target Reads | Mean on-target read length (bp) | Read length N50 (bp) | Covered bases (%) | Mean Sequencing depth (per allele) | Mean sequencing depth (total target region) |
| --- | --- | --- | --- | --- | --- | --- | --- |
| S3 (nCATS-AS) | *CES1P1-CES1* | 59 | 56,822 | 73,710 | 76,011<br>99.94 | 42.76 | 44.42 |
| S3 (nCATS-AS) | *CES1A2-CES1* | 70 | 48,994 | 70,250 | 72,709<br>99.95 | 46.17 |  |
| S3 (nCATS) | *CES1P1-CES1* | 63 | 58,357 | 73,645 | 76,010<br>99.94 | 45.69 | 38.29 |
| S3 (nCATS) | *CES1A2-CES1* | 42 | 53,541 | 70,326 | 71537<br>98.34 | 30.56 |  |

### Misalignment of *CES1A2-*originating reads to *CES1* leads to false-positive SNV calls

*CES1* and *CES1A2* share high sequence similarity, differing primarily in their promoter region, exon 1, and part of intron 1 (Fukami et al. 2008). Traditional short-read sequencing often misclassifies differences in homologous genomic regions, such as *CES1* and *CES1A2* as SNVs. This limitation is well documented, and SNVs within segmental duplications remain systematically under-investigated due to mapping inaccuracies of short reads (Fredman et al. 2004).

Using the multi-target gRNA design strategy for *CES1* enrichment, we observed a subset of reads (with read length N50 = 34.53 kbp) with a high density of SNVs in the region where *CES1* diverges from *CES1A2* when aligned to the *CES1P1-CES1* reference sequence using minimap2 with default parameters (**Fig. 2Ai**). In contrast, realignment of the same reads to our *CES1A2-CES1* custom reference genome markedly reduced mismatches, suggesting that these reads originate from the *CES1A2* locus (**Fig. 2Aii**). This interpretation was further supported by the absence of these SNVs in our end-to-end long-read experiment in the same sample, when competitive mapping was used for sequencing data analysis. By mapping the *CES1A2*-derived reads to the *CES1P1-CES1* reference, we identified 155 SNVs, only 13 of which were also detected in the end-to-end dataset as true *CES1* SNVs (**Fig. 2B**). This indicates that the remaining 142 SNVs are false positives resulting from misalignment of *CES1A2* reads to the *CES1* locus. Interestingly, 89.4% of these incorrect 142 SNVs were associated with unique dbSNP rsIDs in public databases (**Fig. 2B, Supplemental Table S2**), most with very low minor allele frequencies (MAF). This suggests that misalignment artefacts may have contributed to their erroneous inclusion in public variant databases.

**Figure 2.**
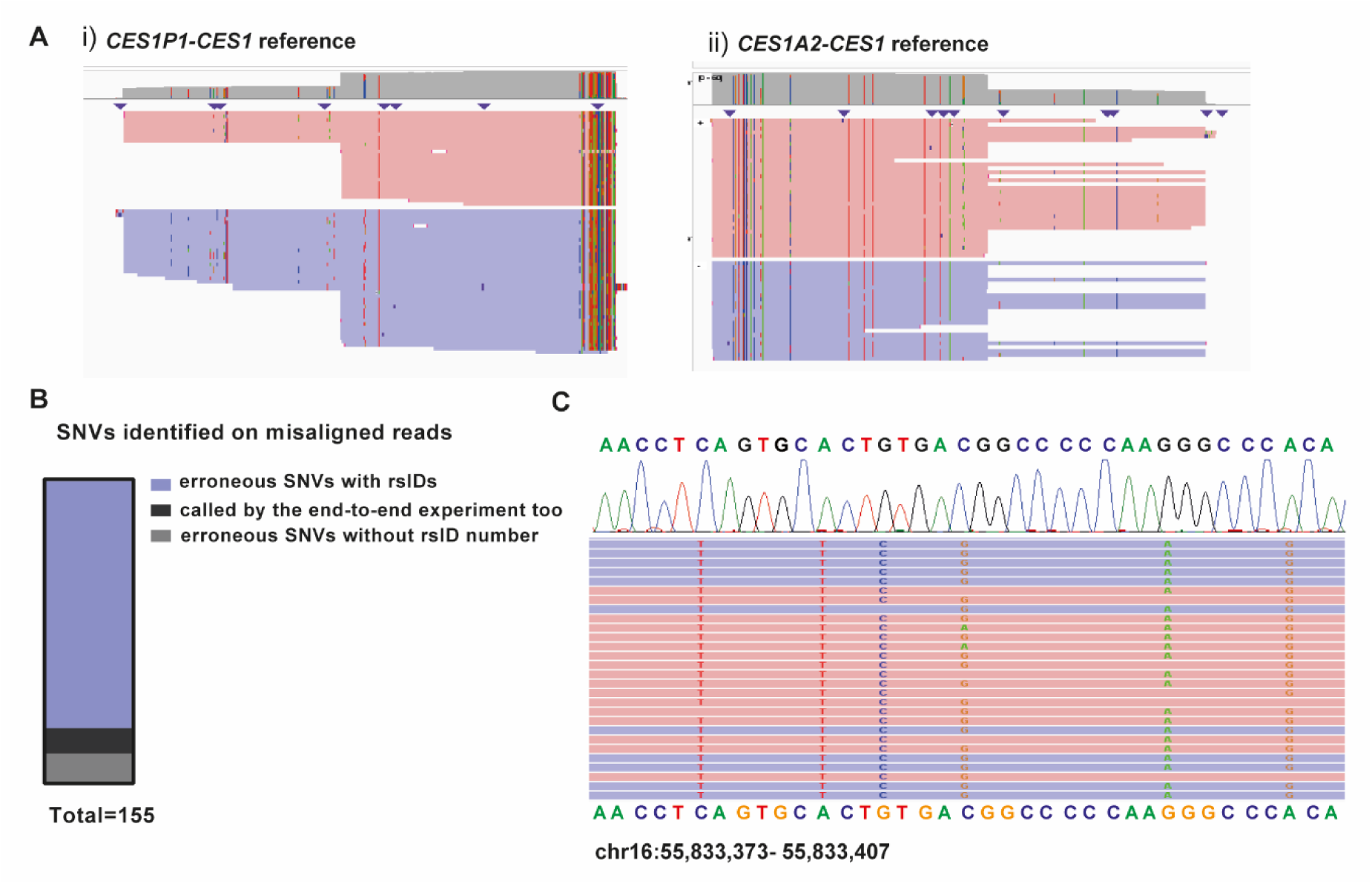
Misclassified *CES1* variants likely originate from *CES1A2-*originating read misalignment. (A) The same set of reads when aligned to (i) *CES1P1-CES1* and (ii) *CES1A2-CES1* reference genome. Red-colored reads indicate alignment to the positive strand, while blue-colored reads indicate alignment to the negative strand. Vertical colored lines within reads represent mismatched bases relative to the reference sequence, and triangles denote insertion events. (B) False positive SNVs called when the *CES1A2*-originating reads are misaligned to *CES1*. (C) Sanger sequencing was done to confirm the absence of the mistakenly called variants on the *CES1* locus and a chromatogram example is depicted.

To test the hypothesis that these SNVs are false positives, we performed PCR and Sanger sequencing of 49 putative erroneous SNV sites in the *CES1* region, 47 of which have assigned rsIDs. PCR primers were designed to anneal to regions containing sequence differences between *CES1* and *CES1A2*, favouring selective amplification of *CES1*. None of these variants were detected in the Sanger chromatograms (**Fig. 2C**), providing strong evidence that they represent misalignment artefacts rather than genuine polymorphisms.

### Robust phasing and variant annotation of the *CES1* locus using nCATS

There is broad recognition that structural variation within the *CES1* locus poses significant challenges for accurate genotyping, leading to limited specificity in conventional *CES1* genotyping methods (Zhu et al. 2012; Ferrero-Miliani et al. 2017; Ikonnikova et al. 2022). Our nCATS approach overcomes these limitations, as long nanopore reads enable variant calling with high mapping accuracy (**Fig. 3A, B**). A total of 764 small variants (SNVs and indels) were called across all analysed samples, 87 of which lacked an rsID (**Fig. 3C**). Of these, 693 variants were identified in the *CES1P1*-carrying haplotype, 221 in the *CES1A2*-carrying haplotype relative to their respective reference sequences, and 150 were shared between both haplotypes. The reliability of SNV detection by nCATS was further validated by Sanger sequencing of *CES1A1a-*, *CES1A1b-*, and *CES1A1c*- specific SNPs, which yielded results consistent with nanopore data calls (**Fig. 3A; Supplemental Table S1**). This may have practical utility in pharmacogenetic diagnostics, particularly in clinical settings where *CES1* genotyping is relevant.

**Figure 3.**
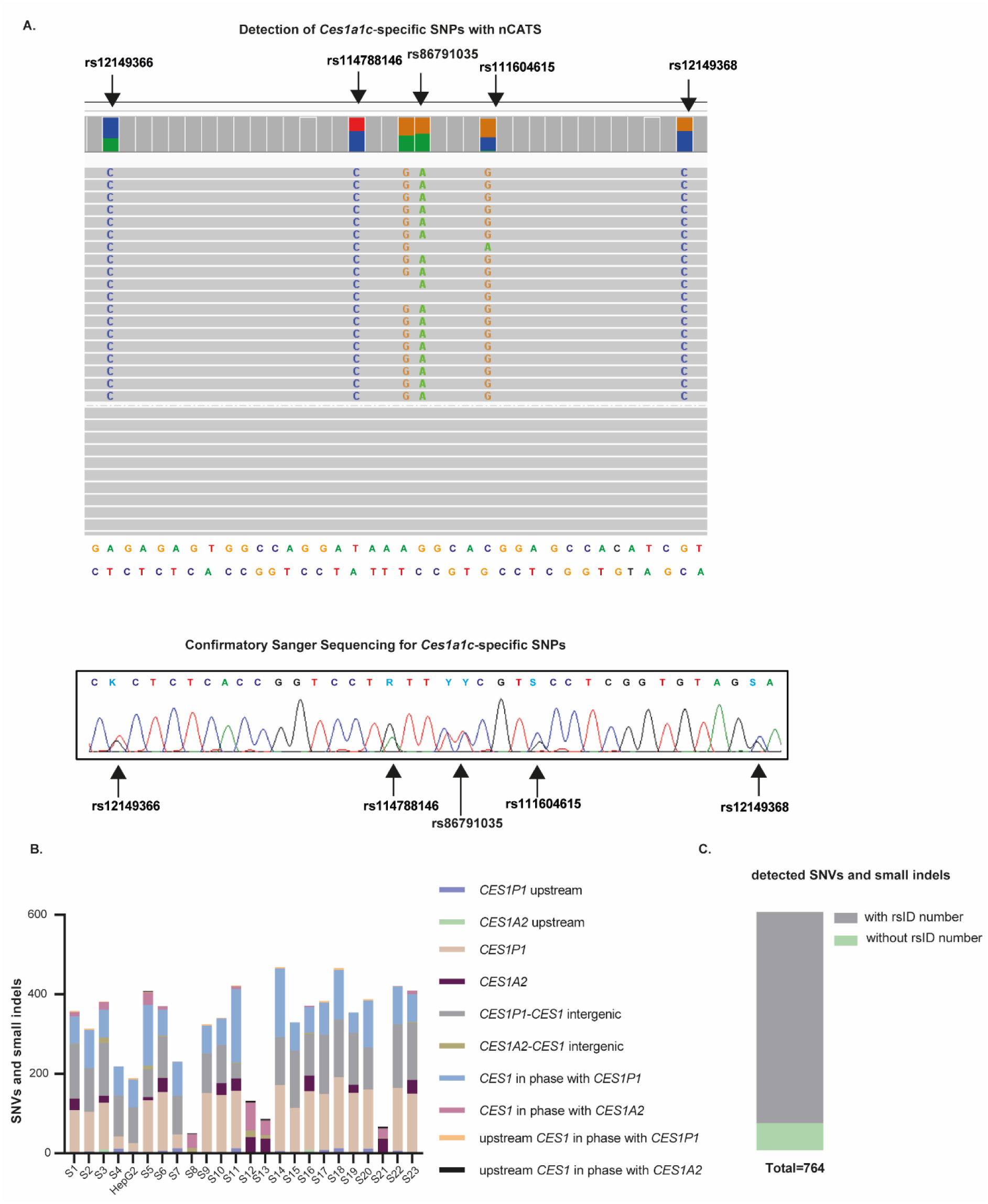
High-resolution SNV profiling of the *CES1* locus using nCATS. (A) *CES1A1c*-specific SNVs as detected by both nCATs and Sanger sequencing in the same DNA sample. (B) Classification of SNVs and small indels by genomic region at the *CES1* locus using nCATS. (C) Summary of the total number of small variants (SNVs and indels) identified across all 24 sequenced samples, indicating how many variants have an rsID and how many are novel (without an rsID).

### Identification of two *CES1P1* subhaplotypes

Phasing *CES1P1–CES1* reads from HepG2 genomic DNA revealed that each allele displayed a distinct SNV pattern within the *CES1P1* locus and its intergenic region with *CES1* (**Fig. 4A**), suggesting the presence of two *CES1P1* subhaplotypes. Comparative analysis showed that all *CES1P1*-carrying alleles could be grouped into one of these two subhaplotypes, here referred to as *CES1P1* type 1 and type 2, based on their SNV profiles (**Fig. 4B, C**). Of the 31 *CES1P1*-carrying alleles analyzed, 25 (80.6%) were of type 1, and 6 (19.4%) were of type 2. Variant analysis using BCFtools identified 32 SNVs shared among both *CES1P1* types, 61 SNVs specific to *CES1P1* type 1, and 144 SNVs specific to *CES1P1* type 2 (**Fig. 4D**). These findings support the existence of two previously uncharacterized *CES1P1* subhaplotypes, each exhibiting distinct sequence signatures that may contribute to regulatory differences.

**Figure 4.**
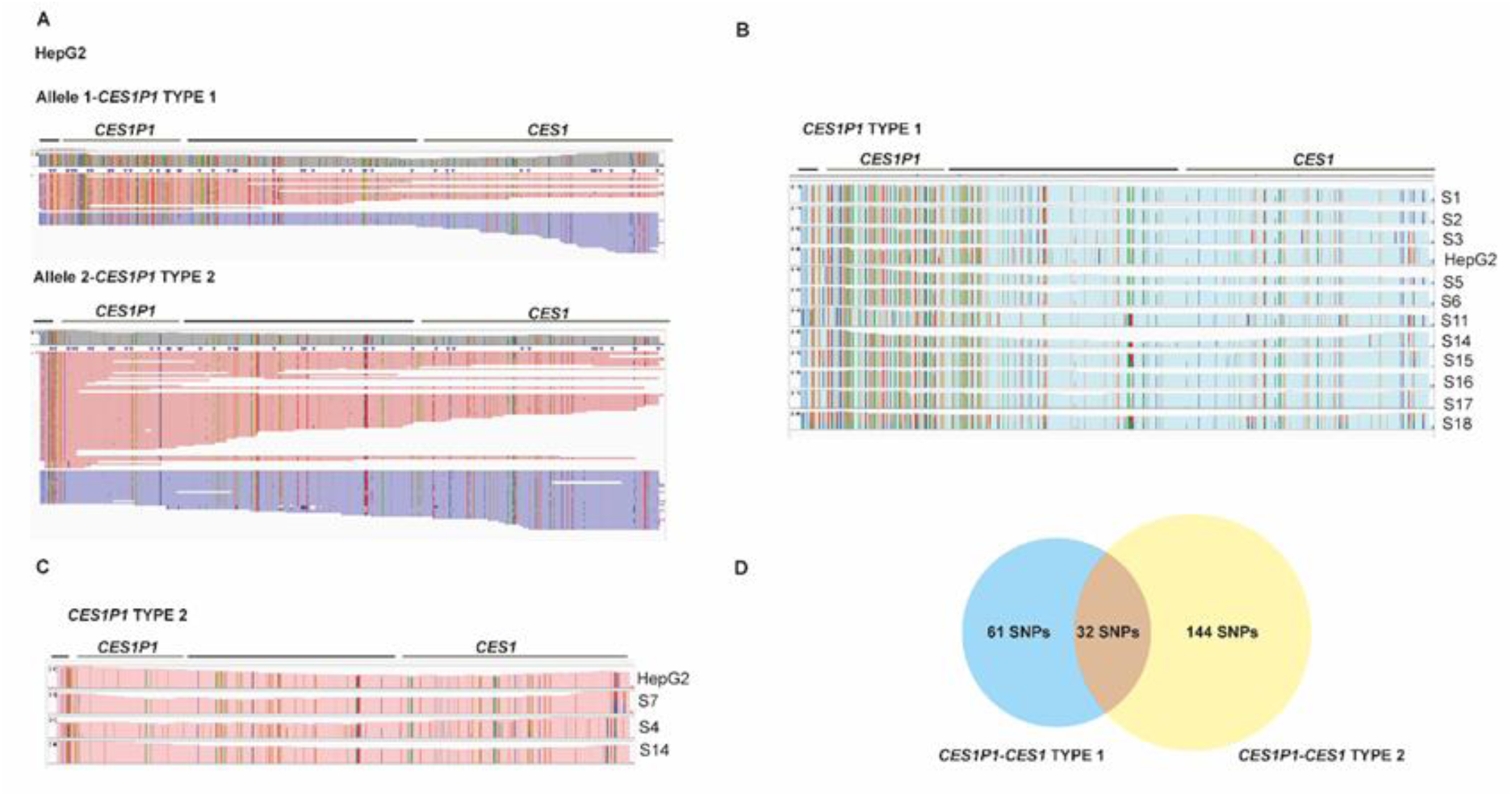
nCATS reveals two distinct *CES1P1* types. (A) IGV view of phased *CES1P1–CES1* reads from HepG2 DNA indicates the presence of two distinct *CES1P1* subhaplotypes. (B) IGV snapshot of nCATS-analyzed alleles carrying *CES1P1* subhaplotype 1. (C) IGV snapshot of nCATS-analyzed alleles carrying *CES1P1* subhaplotype 2. (D) Venn diagram indicating the SNVs (quality score > 20) identified in all analyzed *CES1P1*-carrying alleles (shared across both types) and variants specific to either *CES1P1* type 1 or 2.

### Inverted repeat structures at the *CES1* locus

We also identified an approximately 270-bp DNA rearrangement within the intergenic region between *CES1P1* or *CES1A2* and *CES1*, located approximately 4,480 bp downstream of the *CES1* coding region (**Fig. 5A, B**). This rearrangement was observed with an allele frequency of 43.75% in the analyzed samples. Multiple sequence alignment analysis revealed that this DNA segment is highly conserved among its carriers and represents a reverse complement substitution relative to the reference allele, as evidenced by the high sequence homology between its reverse complement and the reference allele. Moreover, this variant is reported as inversion or partial transposition with respect to hg38 in Human Pangenome Reference Consortium (HPRC) (Liao et al. 2023a) assemblies at chr16:55,798,377-55,798,659, indicating that this structural variant is present in the general population (**Supplemental Fig. S7**).

**Figure 5.**
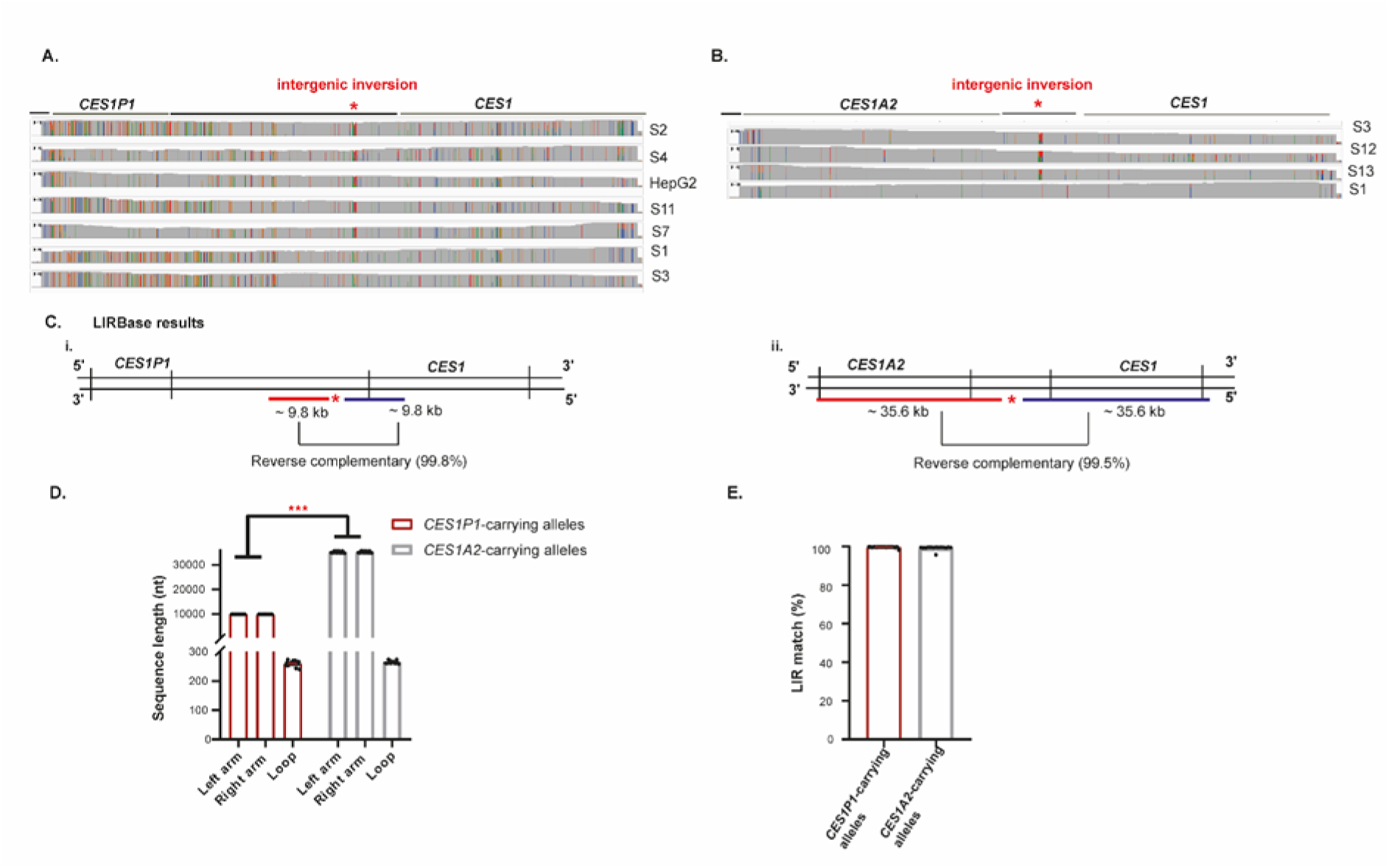
Inverted repeats (IR) on the *CES1* locus. (A) An ≈ 270-bp DNA inversion detected in the *CES1P1-CES1* region (indicated by a red asterisk); this inversion is absent in samples S1 and S3. (B) An ≈ 270-bp DNA inversion detected in the *CES1A2-CES1* intergenic region; this inversion is absent in sample S1. (C) Inverted repeats located within the (i) *CES1P1-CES1* and (ii) *CES1A2-CES1* regions, as annotated by LIRBase. (D) Characteristics of the *CES1* IRs detected by LIRBase, including left and right arm lengths, loop length (E) percentage identity between the arms in *CES1P1*- or *CES1A2*-carrying alleles.

Given the recurrent detection of this inversion, we proceeded to design primers for Sanger sequencing to validate the variant orthogonally. Notably, primers flanking the inversion exhibited bidirectional functionality: forward primers could act as reverse primers and *vice versa*, suggesting the presence of inverted repeats (IRs) in the region. An inverted repeat is a segment of DNA where a sequence is followed, downstream on the same strand by its reverse complement, typically separated by a spacer or loop (Bissler 1998; Lin et al. 2001; Ait Saada et al. 2023). To further investigate this, we used LIRBase, a recently developed database for the identification and annotation of long inverted repeats (Jia et al. 2022).

LIRBase analysis confirmed long inverted repeats (LIRs) flanking the intergenic inversion region in *CES1P1–CES1* alleles, with each arm measuring approximately 9.84 kb and exhibiting 99.85% reverse complementarity (**Fig. 5Ci, D, E**). For *CES1A2–CES1* alleles, each LIR arm spans approximately 35.6 kb, with 99.5% sequence complementarity (**Fig. 5Cii, D, E**).

IRs are known to mediate formation of non-B DNA secondary structures, such as hairpins in singlestranded DNA and cruciforms in double-stranded DNA, via intrastrand annealing (Brázda et al. 2011; Bowater et al. 2022; Burssed et al. 2022). Supporting this, Mfold predictions (Zuker 2003) indicated a strong potential for DNA hairpin formation within the *CES1* IR region (**Supplemental Fig. S8**). The ≈ 270-bp inversion lies precisely within the inverted repeat spacer (or gap), the region flanked by the *CES1* IRs. This spacer likely corresponds to the loop region of a potential IR-mediated hairpin (**Supplemental Fig. S8**), and the presence of genetic variation there results in differential putative hairpin structures (**Supplemental Fig. S9**). DNA or RNA hairpin loops are frequent binding sites for nucleic acid-metabolizing proteins (Nguyen et al. 2022), and genetic variations in the *CES1* loop may impact protein binding affinity and disrupt regulatory function at the *CES1* locus.

In addition, long inverted repeats (LIRs) can be transcribed into long hairpin RNAs (hpRNAs), which may then be processed into small interfering RNAs (siRNAs) (Sasidharan and Gerstein 2008; Watanabe et al. 2008; Ghildiyal and Zamore 2009; Jouravleva and Zamore 2025). A common method to assess the potential of LIRs to produce siRNAs is to align small RNA (sRNA) sequencing data to the genomic sequences of the LIRs of interest. To explore this, we mapped publicly available HepG2 sRNA-seq data (Liao et al. 2023b) to the *CES1* IR genomic sequence. HepG2 cells exhibited small RNA clusters mapping to the *CES1* inverted repeats, with enrichment in the proximal flanking regions of the inverted repeat spacer (**Supplementa**l **Fig. S.10**). This sRNA alignment pattern is consistent with the endogenous siRNA biogenesis pathway, where long RNA hairpins are processed by Dicer into siRNA duplexes.

### Improved characterization of *CES1* haplotype diversity with long-read sequencing and custom *de novo* reference assemblies

To refine the characterization of major *CES1* haplotypes, we leveraged long-read sequencing data generated using our Cas9-targeted nanopore method. Reads > 70 kb in length were phased by individual, and separate *de novo* assemblies were constructed for each allele using the Flye assembler (Kolmogorov et al. 2019). Thirty-seven of the resulting allele contigs were aligned with Clustal Omega and a sequence similarity tree was generated using 1,000 bootstrap replicates. Assemblies were aligned to the *CES1P1-CES1* and *CES1A2-CES1* references and manually inspected in IGV. This approach enabled the identification and reconstruction of the major *CES1* haplotypes, for which we provide updated reference sequences (**Fig. 6A**). We constructed two new reference sequences representing the major *CES1P1–CES1* and *CES1A2–CES1* haplotypes (**Supplemental file S1**), six additional references for *CES1P1–CES1* subhaplotypes as shown in Figure 6A and B, and three new references for *CES1A2– CES1* subhaplotypes as shown in Figure 6A and B (**Supplemental file S2**).

**Figure 6.**
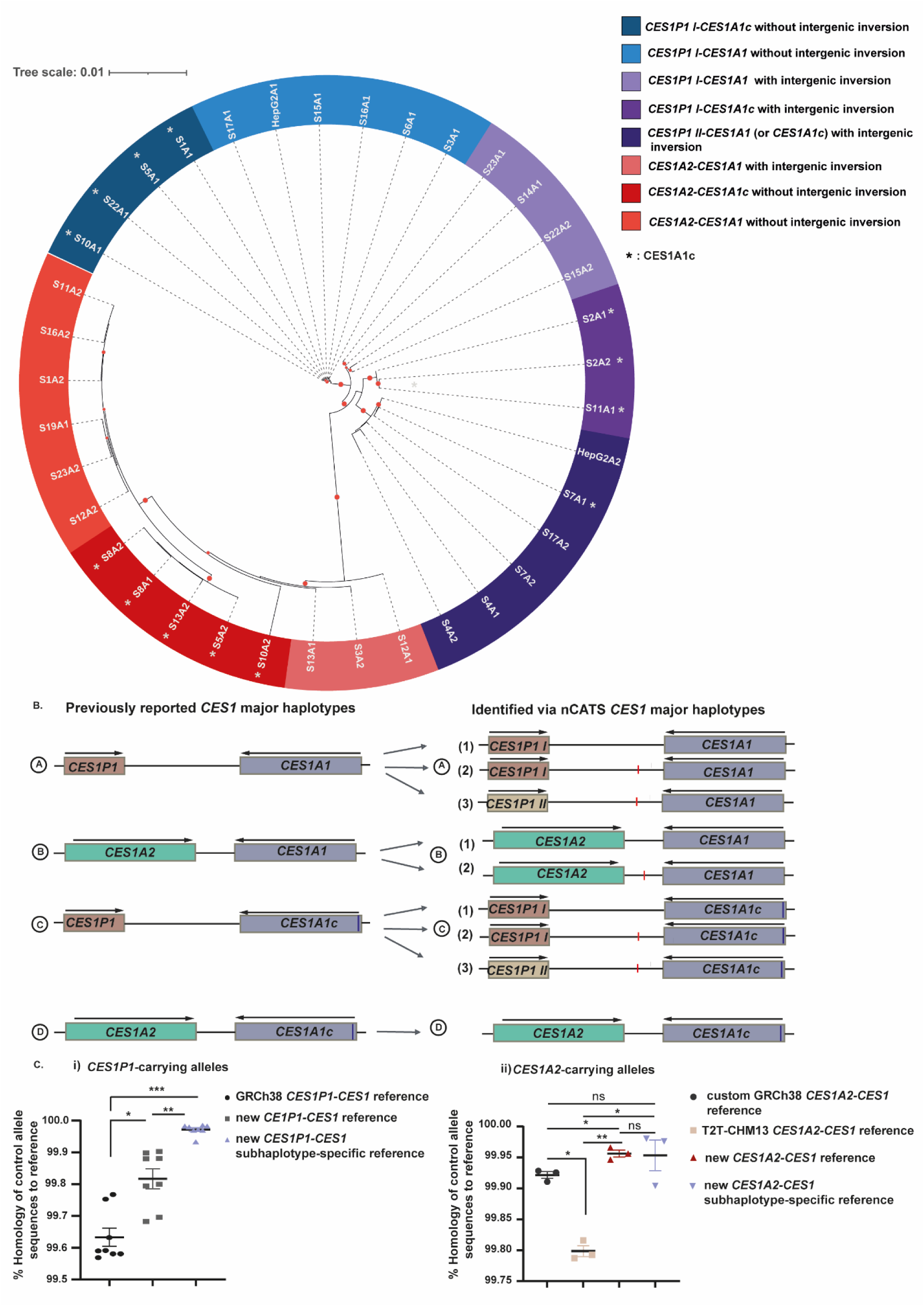
nCATS resolves *CES1* major haplotypes. (A) Sequence similarity tree of allele assemblies from nCATS data. Red circles at branches indicate bootstrap support higher than 90%. A1: allele 1; A2: allele 2 from the corresponding sample. (B) Schematic comparison of previously reported *CES1* major haplotypes with those identified through nCATS (C) Sequence homology of control nCATS-sequenced alleles relative to GRCh38 or T2T-CHM13 references and our new reference sequences. Statistical analysis: paired *t*-test (*P < 0.05; **P < 0.01; ***P < 0.001).

We detected all previously reported major *CES1* haplotypes, characterized by the presence of either *CES1P1* or *CES1A2*, and *CES1A1* or *CES1A1c* within the targeted region (**Fig. 6B**). As expected, haplotypes derived from the *CES1P1-CES1* and *CES1A2-CES1* configurations showed the greatest evolutionary distance to each other, forming distinct clusters in the sequence similarity tree (**Fig. 6A**). The presence of *CES1P1* type 1 or type 2, along with the intergenic inversion, emerged as key determinants of clustering. These structural features enabled the resolution of additional subcategories within the known *CES1* haplotypes (**Fig. 6B**), suggesting the existence of previously uncharacterized genetic diversity that may have important implications for *CES1* regulation and function.

As we performed nCATS in 24 samples (48 alleles) in total, 37 alleles were used to construct the sequence similarity tree and 11 genetically diverse alleles were used as controls to assess how representative the new reference sequences are. We aligned phased reads from these eleven genetically diverse control alleles to both the GRCh38 and T2T-CHM13 reference genomes, as well as to our new haplotype-resolved assemblies. Eight control alleles carried *CES1P1,* representing four distinct *CES1P1–CES1* subhaplotypes (two alleles each of A1, A2, A3, and C2 haplotype, as described in Fig. 6B). We compared the sequence homology of these alleles to the GRCh38.p14 *CES1P1–CES1* reference, our new reference for the major *CES1P1–CES1* haplotype, and subhaplotype-specific *CES1P1–CES1* references. Notably, *CES1P1*-carrying control alleles showed greater homology to our new *CES1P1–CES1* reference sequence than to GRCh38.p14, with further improvement observed when aligned to the corresponding subhaplotype reference (**Fig. 6Ci**).

We next analyzed three *CES1A2*-carrying alleles: two carrying B1 and one D *CES1A2–CES1* subhaplotype (Fig. 6B). These were aligned to: (i) our custom *CES1A2–CES1* GRCh38.p14-based reference, (ii) the *CES1A2–CES1* sequence from the T2T-CHM13v2.0 assembly, (iii) our new reference for the major *CES1A2–CES1* haplotype, and (iv) the corresponding subhaplotype-specific *CES1A2–CES1* references. The T2T-CHM13v2.0 genome was included in this analysis only for *CES1A2*-carrying alleles, as it contains the *CES1A2* gene (albeit annotated as *CES1P1* in the NCBI Genome Data Viewer). Alignment results revealed higher allele sequence homology to our custom GRCh38.p14 *CES1A2–CES1* reference compared with the T2T-CHM13v2.0 reference (**Fig. 6cii**). *CES1A2*-carrying control alleles showed greater identity to our new major *CES1A2–CES1* reference in comparison with both the GRCh38.p14-based and T2T-CHM13v2.0 references (**Fig. 6cii**). In contrast, no substantial increase in homology was observed when aligning to the subhaplotype-specific references (**Fig. 6cii**).

Overall, alignment of *CES1P1*- and *CES1A2-*carrying alleles to our new references consistently resulted in significantly higher sequence identity. These findings highlight the power of long-read sequencing to resolve structurally complex regions and provide more accurate reference sequences for population-scale genomic analyses.

## DISCUSSION

Carboxylesterase 1 (CES1) is a prominent Phase I drug-metabolizing enzyme that participates in the detoxification or activation of numerous therapeutic agents, environmental toxins, as well as endogenous substrates with ester, thioester, or amide bonds (Wang et al. 2018). CES1 accounts for ∼1% of the total liver proteome and contributes to 80–95% of hepatic hydrolytic activity (Imai et al. 2006; Her and Zhu 2020). Despite its importance, the *CES1* locus remains incompletely characterized, due to technical challenges posed by high sequence homology at this genomic region. *CES1* lies tailto-tail with the pseudogene *CES1P1* or the *CES1P1* variant *CES1A2*, an inverted duplication of *CES1* encoding a functional protein. This complex genomic architecture complicates accurate sequencing and variant interpretation, hindering deeper genetic and pharmacogenomic analysis of *CES1*.

Here, we successfully address previous challenges in *CES1* sequencing by employing Cas9-mediated target enrichment combined with long-read nanopore sequencing. This approach generated long ontarget reads spanning up to 76 kb, encompassing the *CES1* pseudogene *CES1P1* (or the variant *CES1A2*), the intergenic region, and the *CES1* prototype gene. Our method overcomes the specificity limitations of conventional *CES1* genotyping assays (Rasmussen and Madsen 2018) and enables highconfidence, haplotype-resolved analysis of *CES1* SVs, and SNVs. Furthermore, our results suggest that certain *CES1* variant entries in public databases may be erroneous, due to potential misalignment artefacts from short-read sequencing.

Importantly, we report some previously unrecognized genetic elements in this region that may expand our understanding of *CES1* function and regulation. Specifically, we identified two distinct *CES1P1* subhaplotypes. Additionally, we detected an approximately 270-nt DNA inversion in the intergenic region between *CES1P1* or *CES1A2* and *CES1*. We found that this inversion variant is flanked by long inverted repeats (LIRs), which can act as genetic elements with a remarkably complex and diverse biological impact. Inverted repeats (IRs) can mediate intra-strand foldback annealing resulting in alternative non-canonical DNA structures, such as hairpins in single-stranded DNA or cruciforms in double-stranded DNA (Brázda et al. 2011; Matos-Rodrigues et al. 2023). In agreement with this, Mfold software predicted DNA hairpin formation in the *CES1* LIR genetic region. These dynamic DNA conformations have been reported to be favoured during biological processes that unwind B-DNA, such as replication or transcription (Wang and Vasquez 2023). IR-induced secondary DNA structures can interfere with cellular processes such as transcription, replication, recombination, and DNA damage and repair (Wang and Vasquez 2023).

IRs are hotspots of genomic instability (Gordenin et al. 1993; Voineagu et al. 2008), as their capacity to form secondary structures can physically hinder DNA polymerase progression, leading to replication fork stalling and DNA double-strand breaks (Lee et al. 2007; Voineagu et al. 2008; Inagaki et al. 2013; Szlachta et al. 2020; Al-Zain et al. 2023). Consistently, IRs are often located near structural variant breakpoints, where they can give rise to diverse structural haplotypes (Vissers et al. 2009; Carvalho et al. 2011; Carvalho and Lupski 2016; Al-Zain et al. 2023; Grochowski et al. 2024). Therefore, we propose that *CES1* LIRs may have played a causal role in the genetic variation and polymorphic evolution of the *CES1* locus. We suggest that instability at the *CES1* IRs may have triggered a DNA double-strand break, which was subsequently resolved through an illegitimate repair pathway, leading to an inverted duplication event and the formation of the *CES1A2–CES1* structural haplotype (**Supplemental Fig. S11**). This model aligns with previous findings demonstrating that inverted repeats facilitate the formation of palindromic duplications (Lin et al. 2001; Tanaka and Yao 2009; Carvalho et al. 2011; AlZain et al. 2023). Although *CES1A2* has already been described as an inverted duplication of *CES1 (Stage et al. 2017a; Stage et al. 2017b)*, our findings indicate that the *CES1A2–CES1* haplotype derives from an inverted duplication of *CES1* plus ≈4.5 kb of its downstream sequence. Moreover, these results are consistent with a previous study showing that chromosome 16, which harbours the *CES1* locus, is enriched in large palindromic duplications (Martin et al. 2004).

Moreover, LIRs can be transcribed into long hairpin RNAs (hpRNAs), which can function as precursors of endogenous small interfering RNAs (siRNAs) (Watanabe et al. 2008; Jia et al. 2022; Jouravleva and Zamore 2025). In line with this, we detected small RNAs mapping to the *CES1* LIRs in the HepG2 hepatic cell line, supporting the hypothesis that *CES1* LIRs might give rise to hairpin-derived endogenous siRNAs. Furthermore, the identification of *CES1* LIRs highlights the importance of using long-read sequencing technologies to accurately characterize the *CES1* locus, as short-read data cannot be unambiguously aligned to long inverted repeat regions.

Finally, we examined major *CES1* haplotypes present in our sequenced alleles by constructing a sequence similarity tree to define the principal clusters. All previously reported *CES1* haplotypes, comprising either *CES1P1* or *CES1A2* and *CES1A1* or *CES1A1c*, were identified. Notably, the presence of the intergenic ≈270-nt inversion variant and/or distinct *CES1P1* subtypes revealed additional sequence clusters, offering deeper insight into the *CES*1 haplotype diversity. Including these newly identified polymorphic elements resulted in a total of nine different haplotypes at the *CES1* locus in the samples analyzed here. We generated new reference sequences for these nine *CES1* haplotypes in addition to two reference sequences for the major *CES1* haplotypes, containing either *CES1P1* or *CES1A2*. These nCATS-derived reference sequences showed higher homology to genetically diverse control alleles across the target region compared with the GRCh38 and T2T-CHM13 reference genomes, supporting the potential of our approach to improve representation of *CES1* haplotype diversity in public databases.

In general, Cas9-targeted nanopore sequencing methods, such as the approach described here, offer a powerful tool to study challenging regions of the human genome. These targeted long-read strategies enable the enrichment of whole target–spanning reads at higher coverage than is typically achievable with whole-genome long-read sequencing. The synergy between targeted long-read strategies and the comprehensive insights provided by the complete telomere-to-telomere human genome assembly (T2T-CHM13) (Nurk et al. 2022), together with emerging human pangenome references (Liao et al. 2023a), creates new opportunities for in-depth genetic and epigenetic analyses. The combination of these resources allows complex or repetitive loci to be studied with a level of accuracy and completeness that was previously impossible.

Overall, this study reveals novel genetic elements within the *CES1* region, laying the groundwork for future research of their biological significance. We anticipate that these findings will support association studies investigating *CES1* polymorphisms in the context of clinical responses to CES1metabolized drugs, thereby enhancing understanding of genetic and phenotypic variability in *CES1* and its relevance for drug safety and effectiveness.

## METHODS

### Patient samples and study approval

This study analyzed 146 anonymized human blood samples (buffy coats) from the Liquid Biobank Bern (https://www.biobankbern.ch/home-en/). The study protocol was approved by the Cantonal Ethics Committee of Bern (BASEC-Nr: Req-2024-00808).

### HepG2 cell culture

The human HepG2 (HB-8065) cell line was obtained from American Type Culture Collection (ATCC). HepG2 cells were maintained in DMEM/F-12 GlutaMAX^TM^ (Gibco) supplemented with 10% of FBS and 1% antibiotics (Penicillin/Streptomycin) and maintained at 37°C and 5% CO2.

### HMW DNA extraction

High molecular weight (HMW) genomic DNA was extracted from human blood samples (n=146), and HepG2 cells using the New England Biolabs Monarch Genomic HMW DNA Extraction Kit for cells and whole blood (NEB #T3010), following the protocol provided by the manufacturer. The concentration and purity of DNA samples were determined with the Qubit ® 4 Fluorometer (Thermo Fisher Scientific, Waltham, MA, USA) and Nanodrop™ One spectrophotometer (Thermo Fisher Scientific, Waltham, MA, USA).

### Guide RNA Design, Cas9 Ribonucleoprotein Complex assembly and cleavage efficiency validation

Custom CRISPR RNAs (crRNAs) were designed using the Integrated DNA Technologies (IDT) design tool and selected based on the highest predicted on-target performance with minimal off-target cuts. CrRNA sequences are provided in **Supplemental Table S3** and gRNA target sites are depicted in **Supplemental Fig. S1**. GnomAD v4.1.0 was used to check for SNV with a minor allele frequency (MAF) > 0.01 at the crRNA binding sites **(Supplemental Table S3).** Functional guide RNAs (gRNAs) were generated by annealing 1 μl of pooled equimolar crRNAs (IDT) with 1 μl of *S. pyogenes* Cas9 transactivating crRNAs (tracrRNAs) (IDT) in 8 μl of duplex buffer (IDT) at 95°C for 5 min followed by cool down at room temperature for 10 min. Cas9-ribonucleoprotein (RNP) complexes were assembled by combining gRNAs (10 μl) with Alt-R® *S. pyogenes* HiFi Cas9 nuclease V3 (IDT) (0.8 μl) in nuclease-free water (79.2 μl) and Reaction Buffer (10 μl) as recommended by Oxford Nanopore Technologies (ONT, SQK-CS9109). The Cas9-RNP cleavage efficiency was initially examined with a PCR-based assay. PCR primers were designed to produce 500-1,000 bp amplicons containing gRNA-target sites (**Supplemental Table S4**), using the Qiagen Multiplex PCR kit. Thermocycler conditions were 95°C for 15 min, 35 cycles of 94°C for 30 s, 58°C for 90 s and 72°C for 60 s, followed by a final extension step of 10 min at 72°C. PCR reactions were purified using ExoSap-IT (Thermo Fisher) and then 3 µl of each amplicon were diluted with 32 µl of nuclease-free H2O before adding 10 µl of assembled Cas9-RNP. Cleavage reactions were incubated at 37°C for 20, 30 or 60 min, then loaded in a 2% agarose TBE gel to resolve the Cas9-cleaved DNA fragments in comparison to the uncut amplicons (**Supplemental Figs. S2-S4**).

### Cas9 enrichment, library preparation and nanopore sequencing with the R9.4.1 flow cell

Libraries for nanopore sequencing were prepared from HMW DNA using the ONT Cas9 sequencing kit (SQK-CS9109) with a modified protocol. The method is schematically summarized in **Fig. 1A**. HMW genomic DNA (30 μl, at a concentration of 200 ng/μl) was dephosphorylated with Phosphatase (4 μl) in the presence of Reaction Buffer (4 μl) for 20 min at 37°C, followed by heating at 80°C for 2 min to deactivate the enzyme. Cas9 RNP complex was prepared as described above. Subsequently, Cas9-RNP complex (10 μl) was added to dephosphorylated gDNA in the presence of Taq polymerase (1 μl) and dATP (1 μl), and incubated for 60 min at 37°C, followed by incubation at 72°C for 10 min. AMX adapters (5 μl) were ligated to phosphorylated DNA ends during a 30 min incubation with T4 ligase (10 μl) in ligation buffer (20 μl) and nuclease-free water (3 μl) at room temperature. The adapter-ligated DNA was cleaned-up with 0.3X AMPure beads (Beckman Coulter), washing twice on a magnetic rack with the long-fragment buffer (250 μl) before eluting, with an elution incubation time of 30 min at 37°C. As DNA libraries were frequently very viscous, samples were centrifuged at maximum speed for 3 min before applying the magnet at the elution step. Sequencing libraries were prepared by mixing 12 μl of DNA library with 37.5 μl sequencing buffer and 25.5 μl loading beads. Long read sequencing was performed on a GridION X5 device (ONT) equipped with R9.4.1 flow cells (ONT) for 24 h. The run was paused 30 min after its initiation and flow cell was loaded with additional library material to increase the number of total reads. In one of our runs, we employed adaptive sampling, and the reference genomes used for read alignment are listed in **Supplemental Table S5.**

### Library preparation adjustment to make it compatible with the R10.4.1 flow cell

As the ONT Cas9 sequencing kit (SQK-CS9109) was discontinued, we replaced the following reagents: Reaction Buffer from the SQK-CS9109 kit was replaced with rCutSmart™ Buffer (B6004S, New England Biolabs), dATP Solution (N0440S, New England Biolabs) and Taq DNA Polymerase (M0273S, New England Biolabs) were used for dA-tailing. The adapter ligation step was carried out with Salt-T4® DNA Ligase (M0467L, New England Biolabs) and Ligation Sequencing Kit V14 (SQK-LSK114, Oxford Nanopore Technologies). The nanopore Cas9-targeted sequencing (nCATS) protocol was kept unchanged until the cleavage and dA-tailing. Subsequent steps, including DNA clean-up, adapter ligation, second DNA clean-up, final library elution, and loading, were performed according to the SQKLSK114 protocol. Libraries were loaded into R10.4.1 flow cells and sequenced for 24 h. The sequencing run was paused 30 min after sequencing initiation to reload additional library material, as routinely performed for R9.4.1 flow cells.

### Analysis of Oxford Nanopore sequencing data

Our bioinformatics pipeline is summarized in **Supplemental Fig. S12**. Live basecalling was performed with Guppy (high-accuracy model, v4.3.0), and reads with a mean quality score <9 were discarded. Reads shorter than 3 kb were removed with SAMtools v1.22.1 (Danecek et al. 2021). Filtered reads were competitively mapped with minimap2 (v2.29)(Li 2018) to a two-contig haplotype reference containing sequences for the two major *CES1* structural haplotypes (*CES1P1–CES1* and *CES1A2–CES1*) with the --preset lr:hq parameter. The *CES1P1-CES1* reference sequence corresponds to chr16:55,758,219-55,834,270 of the GRCh38.p14 primary assembly. The *CES1A2-CES1* reference was constructed by combining the AB119998.1 sequence of the *CES1A2* gene (https://www.ncbi.nlm.nih.gov/nuccore/AB119998.1) (Hosokawa et al. 2008) with the flanking chr16:55,758,219-55,760,576 upstream sequence and chr16:55,794,158-55,834,270 downstream sequence from the GRCh38.p14 primary assembly, as reported for the human gene *CES1P1* (ENST00000702195.1) from the GENCODE V48 *CES1A2* position. Alignments with a dynamic programming (DP) score>1000 were retained. BAM files were filtered and reads with a mapping quality lower than 20 were removed with SAMtools (v1.22.1). To retain only primary mapped reads, we filtered the alignment files with SAMtools (v1.22.1) using the -F 260 parameter, which excludes unmapped and secondary alignments. On-target alignments were visualized with the Integrative Genomics Viewer (IGV) (v2.16)(Thorvaldsdóttir et al. 2013). Sequencing depth and coverage were computed with Mosdepth (v0.3.10)(Pedersen and Quinlan 2018) and PanDepth (v2.25)(Yu et al. 2024). Depth and distribution of read alignments to each major haplotype-specific reference were used to infer the structural variant genotype for each sample. For *CES1P1/CES1A2* heterozygous samples, the final BAM file was split by reference with SAMtools (v1.22.1), and per-haplotype FASTQ files were extracted using SAMtools (v1.22.1). Alignment quality metrics were assessed on the extracted FASTQ files with NanoPlot (v1.46.1)(De Coster and Rademakers 2023). Small variant calling was performed with Clair3 (v1.2.0)(Zheng et al. 2022), using the corresponding haplotype reference (*CES1P1–CES1* or *CES1A2–CES1*). VCF files were filtered to retain only high-quality calls (q>20) with awk (awk ’$1 ∼ /^#/ || $6 > 20’ input.vcf > hqinput.vcf). Filtered VCFs were processed by a custom rsID retrieval tool (https://github.com/ayouballah/rsID_retrieval) to annotate variants with rsIDs from the NCBI variations and Entrez database. For homozygous samples for either the *CES1P1* or *CES1A2-* containing haplotype, reads were realigned to the corresponding single-contig reference with minimap2 (v2.29) and all downstream steps were performed as described above.

### Alignment of high-throughput sRNA sequencing data to *CES1* long inverted repeats (LIR)

Small-RNA (sRNA) sequencing data for the HepG2 cell line were downloaded from the NCBI Sequence Read Archive (Liao et al. 2023b) (NCBI BioProject database, accession number PRJNA889826). Small RNA read quality was assessed with FastQC (v.0.74)(Andrews 2010). Reads were mapped to the *CES1P1-CES1* reference genome using Bowtie (v2.5.3)(Langmead and Salzberg 2012) with no mismatch allowed. The *CES1P1-CES1* reference was used, as our sequencing showed HepG2 is homozygous for *CES1P1*.

### Sequence similarity tree construction and major *CES1* haplotype identification

Variant calling was performed with Clair3 (v1.2.0) and haplotype phasing was carried out using WhatsHap phase (v2.8)(Martin et al. 2023). The phased variants were then used to tag the mapped reads using WhatsHap haplotag (v2.8) and WhatsHap split (v2.8) was used to partition the raw reads by haplotype, resulting in two haplotype-specific read sets for each sample. Reads were filtered to retain reads with lengths longer than 70 kb. For sample S9, haplotype phasing was improved by manual curation. For CES1P1/CES1A2 heterozygous samples, the reads were already phased during the step in which the BAM file was split by reference using SAMtools (v1.22.1). The phased reads were assembled *de novo* into haplotype-resolved contigs using Flye (v2.9.6)(Kolmogorov et al. 2019) (**Supplemental Fig.S12**). Assembled contig orientation was assessed following alignment to the reference sequence, and all assemblies were converted to the positive strand. Input reads were mapped back to the assembly using minimap2 (v2.29) with a dynamic programming (DP) score>1000 to assess assembly quality. Thirty-seven allele contigs (23 *CES1P1*-carriers and 14 *CES1A2*-carriers) were merged and subsequently aligned using Clustal Omega (v1.2.4)(Sievers et al. 2011), with the output generated in PHYLIP format. To infer sequence similarity, a maximum likelihood tree was constructed using RAxML (v8.2.13)(Stamatakis 2014) with 1,000 bootstrap replicates. The final phylogenetic tree was visualized using the Interactive Tree of Life (iTOL) in circular mode. Major *CES1* haplotypes were determined, and de novo assembly was performed using Flye (v2.9.6) for each major haplotype, using the allele contigs constituting each *CES1* haplotype. New reference sequences are provided in **Supplementary files 1,2**.

#### Evaluation of new reference sequences

We assessed the accuracy and representativeness of the newly generated reference sequences using 11 genetically diverse alleles obtained via nCATS long-read sequencing: eight *CES1P1*-carrying alleles and three *CES1A2*-carrying alleles. These alleles were not included in the initial sequence similarity tree, which we used to construct the new references. The selection of these control alleles was based on their genetic diversity, allowing us to assess how well the new reference sequences represent the diversity of *CES1* haplotypes. For *CES1P1*-carrying alleles, we mapped phased reads to (i) the *CES1P1– CES1* reference sequence from the GRCh38.p14 primary assembly (chr16: 55,758,219–55,834,270), (ii) the newly constructed major *CES1P1–CES1* reference, and (iii) the corresponding set of *CES1P1–CES1* subhaplotype reference sequences. For *CES1A2-*carrying alleles, we aligned phased reads against (i) the custom *CES1A2–CES1* reference from the GRCh38.p14 primary assembly, (ii) the *CES1A2–CES1* cleavage product reference extracted from the T2T-CHM13 assembly (chr16:61,556,459–61,629,268), (iii) the newly constructed *CES1A2–CES1* major haplotype reference, and (iv) the *CES1A2–CES1* subhaplotype reference sequences. The T2T-CHM13-derived reference was only used for *CES1A2* alleles, as it includes the *CES1A2* gene but not *CES1P1*. Alignments of phased reads were performed using a dynamic programming alignment strategy with alignment scores exceeding 1000 with minimap2 (v2.29). Resulting BAM files were filtered to retain only alignments with mapping quality (MAPQ) >20. Variant calling was conducted using Clair3 (v.2.0). Sequence homology was calculated as: Sequence Homology = 1 − (number of mismatches / reference length).

### Sanger Sequencing

PCR amplification for *CES1A1a-c* identification was performed with the QIAGEN Multiplex PCR Kit™, using the following primers: CES1_Promoter_F:5’-GCT CTA ACA TTT TCC AGT TGT T-3’ and CES1_Promoter_R: 5’-TCC AAG TCC TAA TAT GGA AGT CGT G-3’, as previously described (Yamada et al. 2010) (Hamzic et al. 2017). Primers for screening the erroneous *CES1* SNPs described in Fig.2 are listed in **Supplemental Table S6**. Exonuclease I - Shrimp alkaline phosphatase cleaning was performed with the PCR products. The sequencing reactions were done with the BigDye Terminator v3.1 Cycle Sequencing Kit (Thermo Fisher Scientific). After cleanup with the BigDye XTerminator Purification Kit (Applied Biosystems), samples were sequenced on a 3130xl™ Genetic Analyzer. Chromatograms were visualized with Sequencher® 5.1.

### Multiplex long-range CES1P1/CES1A2 allele-specific PCR

Allele-specific PCR was used to genotype for *CES1P1/CES1A2* (**Supplemental Fig. S5**). The following primers were used: CES1P1_F: 5’-GCG ACC TCC GTA CTT GGA A-3’, CES1P1_R: 5’-GTT CAA GAC AGA GGA ATG CAT GA-3’ and CES1_R: 5’-GGA CTG TGA GGG TAC ATA CGG-3’ (Bjerre et al. 2018) to amplify specific segments from either *CES1P1* or *CES1A2* with PrimeSTAR® GXL DNA Polymerase (Takara). Thermocycler conditions were: initial denaturation at 94°C for 2 min, 35 cycles of denaturation at 98°C for 10 s, annealing at 58°C for 15 s and extension at 68°C for 10 min, followed by final extension at 68°C for 10 min. PCR products were separated and visualized on 0.8% agarose gels.

## DATA ACCESS

Raw sequencing data generated in this study are available to reviewers at the following temporary link: https://cloud.bioinformatics.unibe.ch/index.php/s/AbGcpcBs69Jirsi. The data will be deposited in a public repository and assigned permanent accession numbers prior to publication.

## AKNOWLEDGMENTS

This research was supported by the Interfaculty Bioinformatics Unit of the University of Bern under the Margaret Dayhoff Centenary Grant. We thank Sonja Gempeler and Dr. Loïc Borcard for their help during Nanopore sequencing (both at the Institute for Infectious Diseases, University of Bern). The Nanopore sequencing data were generated in collaboration with the IFIK NGS platform, Bern, Switzerland. This study was funded by Swiss National Science Foundation (SNSF), grant number 320030_212583.

## Author contributions

C.R.L designed and supervised the study. E.L. designed, performed and analyzed experiments and interpreted results with C.R.L., U.A., and A.R. A.A., A.N. and A.B. contributed to the development of the data analysis pipeline used in this study. E.L. wrote the manuscript. All authors discussed the results and commented on the manuscript.

